# Evolutionary Stalling and a Limit on the Power of Natural Selection to Improve a Cellular Module

**DOI:** 10.1101/850644

**Authors:** Sandeep Venkataram, Ross Monasky, Shohreh H Sikaroodi, Sergey Kryazhimskiy, Betül Kaçar

## Abstract

Cells consist of molecular modules which perform vital biological functions. Cellular modules are key units of adaptive evolution because organismal fitness depends on their performance. Theory shows that in rapidly evolving populations, such as those of many microbes, adaptation is driven primarily by common beneficial mutations with large effects, while other mutations behave as if they are effectively neutral. As a consequence, if a module can be improved only by rare and/or weak beneficial mutations, its adaptive evolution would stall. However, such evolutionary stalling has not been empirically demonstrated, and it is unclear to what extent stalling may limit the power of natural selection to improve modules. Here, we empirically characterize how natural selection improves the translation machinery (TM), an essential cellular module. We experimentally evolved populations of *Escherichia coli* with genetically perturbed TMs for 1,000 generations. Populations with severe TM defects initially adapted via mutations in the TM, but TM adaptation stalled within about 300 generations. We estimate that the genetic load in our populations incurred by residual TM defects ranges from 0.5 to 19%. Finally, we found evidence that both epistasis and the depletion of the pool of beneficial mutations contributed to evolutionary stalling. Our results suggest that cellular modules may not be fully optimized by natural selection despite the availability of adaptive mutations.

## Introduction

Biological systems are organized hierarchically, from molecules to cells, organisms and populations (1–5). At the lowest level, macromolecules form cellular modules, such as the translation machinery, or other metabolic pathways (4, 6–9). Different modules perform different cellular functions, which together determine the fitness of the organism. Adaptive mutations improve these modules, enabling organisms to consume new resources (10, 11), become resistant to drugs (12–14), thrive in otherwise harsh conditions (15–17), etc. However, not all beneficial mutations have the same chance of becoming fixed in populations. The evolutionary dynamics that govern the fates of mutations can be very complex, particularly in large populations with limited recombination (14, 18–23). Recent theoretical models and experimental observations characterized these dynamics at the genetic and fitness levels (18, 21, 22, 24–27). However, our understanding of these dynamics at the level of cellular modules and their implications for the evolution of physiological functions remains poor (28).

When beneficial mutations are rare (i.e., if the population evolves in the successional mutations regime (24)), or when recombination rates are high, modules evolve independently of each other, so that the speed of adaptation of each module depends only on the supply and the fitness effects of mutations in that module alone (29). Natural selection can only improve a module if there are mutations available to improve it (that is, the module is not at a local performance peak) and if their fitness benefits are above ∼1/*N*, the inverse of the population size (30–34). Module adaptation may also stop before reaching this threshold, when a balance is reached between fixations of mutations that improve the module and those that degrade it (31, 35–37). Regardless of the reasons for why some modules are not improvable, in this regime, all improvable modules keep adapting simultaneously (albeit at possibly different rates).

In many populations, particularly in microbes, beneficial mutations are common and recombination is rare, so that the evolutionary fates of different adaptive mutations are not independent (14, 19, 20, 23). In this so-called concurrent mutations regime, adaptation is primarily driven by mutations with large fitness effects, whereas the evolutionary fates of mutations whose effects fall below a certain “emergent neutrality threshold” are largely determined by the genetic backgrounds in which they arise (26, 38, 39). The emergent neutrality threshold depends on the supply and the fitness effects of all adaptive mutations in the genome. Thus, the rate of adaptation in any one module depends not only on the mutations in that module but also on mutations in all other modules, through the emergent neutrality threshold.

Coupling of modules by the emergent neutrality threshold has important implications for the evolutionary dynamics of individual modules. In particular, improvable modules accumulate adaptive mutations only if such mutations arise frequently and provide fitness benefits above the emergent neutrality threshold. In contrast, modules that are improvable only by rare mutations, or mutations whose fitness benefits are too small, will not adapt (29). We refer to a failure of an otherwise improvable module to accumulate adaptive mutations as “evolutionary stalling”.

Evolutionary stalling can theoretically limit the power of natural selection to improve a module, but whether stalling occurs in biological systems is unclear. There are plausible scenarios in which evolutionary stalling would not occur. For example, if many modules are improvable by mutations with similar rates and fitness effects, all of them would improve simultaneously, without exhibiting evolutionary stalling. Some evolution experiments appear to support this possibility (18, 40–44). However, in some conditions, natural selection focuses on improving a single or a handful of modules (41, 43, 45–54). For example, resistance mutations in bacteria often occur in the protein complex whose function is inhibited by an antibiotic (13, 55, 56). Similarly, compensatory mutations after a severe genetic perturbation are often concentrated in a handful of pathways with functional relationships to the perturbation (41, 43, 45, 49, 50, 53, 54). The fact that adaptive mutations are not observed in some other modules can be interpreted in two ways. Either these modules exhibit evolutionary stalling or they are not improvable in the first place. Previous studies have not attempted to distinguish between these two possibilities.

Whenever there are modules whose adaptation is stalled, we would generically expect the focus of natural selection to shift over time from some modules to others, even in a constant selective environment. Such shifts should occur because the emergent neutrality threshold that determines which modules adapt and which ones stall is a dynamic quantity. As mentioned above, this threshold depends on the supply and the effects of all adaptive mutations in the genome, which themselves change over time by at least two genetic mechanisms. First, as a population adapts, the supply of available adaptive mutations is being depleted, with large-effect mutations being depleted first (57). Second, genetic interactions (epistasis) can modulate the fitness benefits of mutations (40, 58–60) or even open up and close down large pools of adaptive mutations (10, 41, 61, 62). A recent study of a 60,000-generation long evolution experiment in *Escherichia coli* found evidence that both of these genetic mechanisms cause temporal shifts in the statistical distribution of mutations among genes and operons (57). However, this study did not identify the specific modules in which adaptive evolution stalled or resumed, nor did it quantify whether evolutionary stalling imposes any limit on the power of selection to improve any specific module.

Here, we use experimental populations of the bacterium *Escherichia coli* (*E. coli*) to explicitly demonstrate the onset of stalling in the evolution of the translation machinery and to quantify the fitness cost imposed by it. In *E. coli*, the core macro-molecular complexes that carry out translation are encoded by a well annotated set of about 200 genes (8, 9, 63). We operationally define this set as the translation machinery module, or TM for short. To detect evolutionary stalling, we first disrupt the TM by replacing the native Elongation Factor Tu (EF-Tu) in *E. coli* with several of its orthologs (64, 65). We then evolve these strains in rich media where rapid and accurate translation is required for fast growth (66). We thus expect natural selection to favor adaptive mutations in the TM. In addition to such TM-specific mutations, mutations that improve the performance of other cellular modules could also be adaptive. We refer to such mutations as “generic”. We show the TM-specific and generic mutations compete against each other in our populations. We then directly demonstrate that TM adaptation stalls in at least some populations and estimate the genetic load caused by stalling. Finally, we characterize the genetic mechanisms that contribute to evolutionary stalling in our populations.

## Results

We previously replaced the native EF-Tu in *E. coli* with its orthologs from *Salmonella typhimurium, Yersinia enterocolitica, Vibrio cholerae* and *Pseudomonas aeruginosa* and one reconstructed ancestral variant (64) (Table 1, Dataset S1). EF-Tu is encoded in *E. coli* by two paralogous genes, *tufA* and *tufB*, with the majority of the EF-Tu molecules being expressed from *tufA* (67). To replace all EF-Tu molecules in the cell, the *tufB* gene was deleted and the foreign orthologs were integrated into the *tufA* locus (64). We also included the control strain in which the *tufB* gene was deleted and the original *E. coli tufA* was left intact. We refer to the engineered “founder” *E. coli* strains as E, S, Y, V, A and P by the first letter of the origin of their *tuf* genes (Table 1).

**Table 1.** Founders used for the evolution experiment.

| Strain | EF-Tu origin species | Number of amino acid differences from *E. coli* EF-Tu (percent identity) | Fitness relative to E strain ± SEM, % per generation |
| --- | --- | --- | --- |
| E | *Escherichia coli* (control) | 0 (100) | $0 \pm 0.4$ |
| S | *Salmonella typhimurium* | 1 (99.75) | $+0.49 \pm 0.05$ |
| Y | *Yersinia enterocolitica* | 24 (93.91) | $-3.02 \pm 0.02$ |
| V | *Vibrio cholerae* | 51 (87.06) | $-19.0 \pm 0.7$ |
| A | Reconstructed ancestor | 21 (94.67) | $-34.4 \pm 0.4$ |
| P | *Pseudomonas aeruginosa* | 62 (84.38) | $-35.0 \pm 0.1$ |

We first quantified the TM defects in our founder strains (Methods). Kaçar et al. showed that EF-Tu replacements lead to declines in the *E. coli* protein synthesis rate and proportional losses in growth rate in the rich laboratory medium LB (64). In our subsequent evolution experiment, natural selection will favor genotypes with higher competitive fitness, which may have other components in addition to growth rate (68–72). We confirmed that EF-Tu replacements caused changes in competitive fitness relative to the control E strain (Table 1), and that competitive fitness and growth rate were highly correlated (SI Appendix, Fig. S1). We conclude that the competitive fitness of our founders in our environment reflects their TM performance. Further, we found that the fitness of the S and Y founders were similar to that of the control E strain (≤ 3% fitness change) indicating that their TMs were not substantially perturbed. In contrast, the fitness of the V, A and P founders were dramatically lower (≥ 19% fitness decline; Table 1) indicating that their TMs were severely perturbed. Note that the E strain itself carries an 4% fitness cost compared to the wildtype *E. coli* strain (Methods, Dataset S2).

### Clonal interference slows down TM adaptation

We instantiated 10 replicate populations from each of our six founders (60 populations total) and evolved them in LB for 1,000 generations with daily 1:10^4^ dilutions and the bottleneck population size *N* = 5×10^5^ cells (Methods). The competitive fitness of all but one of the populations against their respective founders significantly increased during evolution (*P* < 0.05, t-test after Benjamini-Hochberg correction; Fig. 1A, Dataset S3), with significantly larger increases observed in the V, A, and P populations compared to E, S, and Y populations (ANOVA, *P* < 10^−16^). Furthermore, 24 out of 30 E, S, and Y populations and 3 of the 30 V, A and P populations became more fit than the control E strain with 95% confidence (Fig. 1B, Dataset S4).

**Figure 1.**
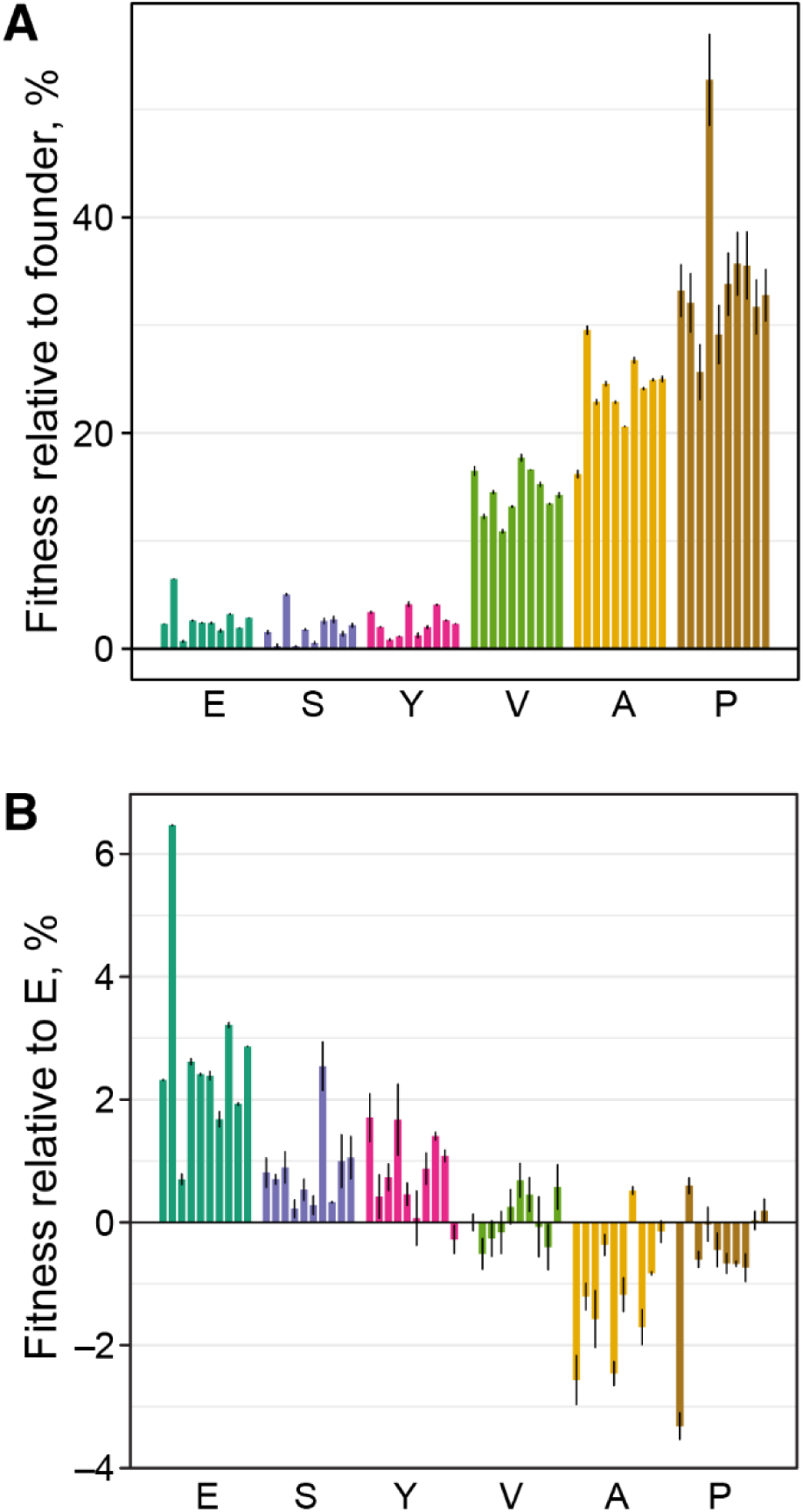
Competitive fitness of evolved populations. **A**. The competitive fitness of evolved populations relative to their respective founders. **B**. The competitive fitness of evolved populations relative to the E strain. For both panels, fitness is measured in % per generation, and error bars show ±1 SEM (see Datasets S3, S4).

Adaptation in these populations can be driven not only by mutations in the TM, but also by mutations in other modules. However, evolutionary stalling in the TM can occur only if mutations improving the TM compete against other types of mutations within the same population. To determine whether both types of mutations occur in our populations, we conducted whole-population whole-genome sequencing at multiple timepoints throughout the evolution experiment. This sequencing strategy allows us to directly observe competition dynamics between mutations in different modules (18, 57, 73, 74).

We selected replicate populations 1 through 6 descended from each founder (a total of 36 populations), sampled each of them at 100-generation intervals (a total of 11 time points per population) and sequenced the total genomic DNA extracted from these samples. We developed a bioinformatics pipeline to identify *de novo* SNPs and small insertion/deletion events in this data set (Methods). Then, we called a mutation adaptive if it satisfied two criteria: (i) its frequency changed by more than 20% in a population; and (ii) it occurred in a “multi-hit” gene, i.e., a gene in which two independent mutations passed the first criterion. We augmented our pipeline with the manual identification of large-scale copy-number variants which could only be reliably detected after they reached high frequency in a population (Methods and SI Appendix, Fig. S2). We successfully validated 43 / 45 tested variants via Sanger sequencing (Methods, Dataset S5)

We first identified and broadly classified the targets of adaptation and documented their competition dynamics across all populations, irrespectively of their founder. Our filtering procedure yielded a total of 167 new putatively adaptive mutations in 28 multi-hit genes, with the expected false discovery rate of 13.6%, along with an additional 11 manually-identified chromosomal amplifications, all of which span the *tufA* locus (Methods, Dataset S6 and SI Appendix, Figs. S2, S3). We classified putatively adaptive mutations as “TM-specific” if the genes where they occurred are annotated as translation-related (Methods). We classified all other mutations as “generic”. We found that 38 out of 178 (21%) putatively adaptive mutations in 6 out of 28 multi-hit genes were TM-specific (Dataset S6). This is significantly more than expected by chance (*P* < 10^−4^, randomization test) since the 215 genes annotated as translation-related comprise only 4.3% of the *E. coli* genome. All of the TM-specific mutations occurred in genes whose only known function is translation-related, such as *rpsF* and *rpsG*, suggesting these mutations arose in response to the initial defects in the TM. The set of TM-specific mutations is robust with respect to our filtering criteria (SI Appendix, Fig. S4).

TM-specific mutations occurred in 17 out of 36 sequenced populations. Generic mutations were also observed in all of these populations (SI Appendix, Fig. S3). Thus, whenever TM-specific mutations occurred, generic mutations also occurred, such that the fate of TM-specific mutations likely depended on the outcome of competition between mutations within and between modules (Fig. 2). As a result of this competition, only 14 out of 27 (52%) TM-specific mutations that arose (excluding 11 *tufA* ampliciations) went to fixation, while the remaining 13 (48%) succumbed to clonal interference (Fig. 2 and SI Appendix, Fig. S3). In at least two of these 13 cases, a TM-specific mutation was outcompeted by expanding clones driven by generic mutations: in population V6, a TM-specific mutation in *fusA* was outcompeted by a clone carrying generic mutations in *fimD, ftsI* and *hslO* (Fig. 2); and in population P3, a TM-specific mutation in *tufA* was outcompeted by a clone carrying generic mutations in *amiC* and *trkH* (Fig. 2). We conclude that, while TM-specific beneficial mutations are sufficiently common and their fitness effects are at least sometimes large enough to successfully compete against generic mutations, clonal interference reduces the power of natural selection to recover TM performance.

**Figure 2.**
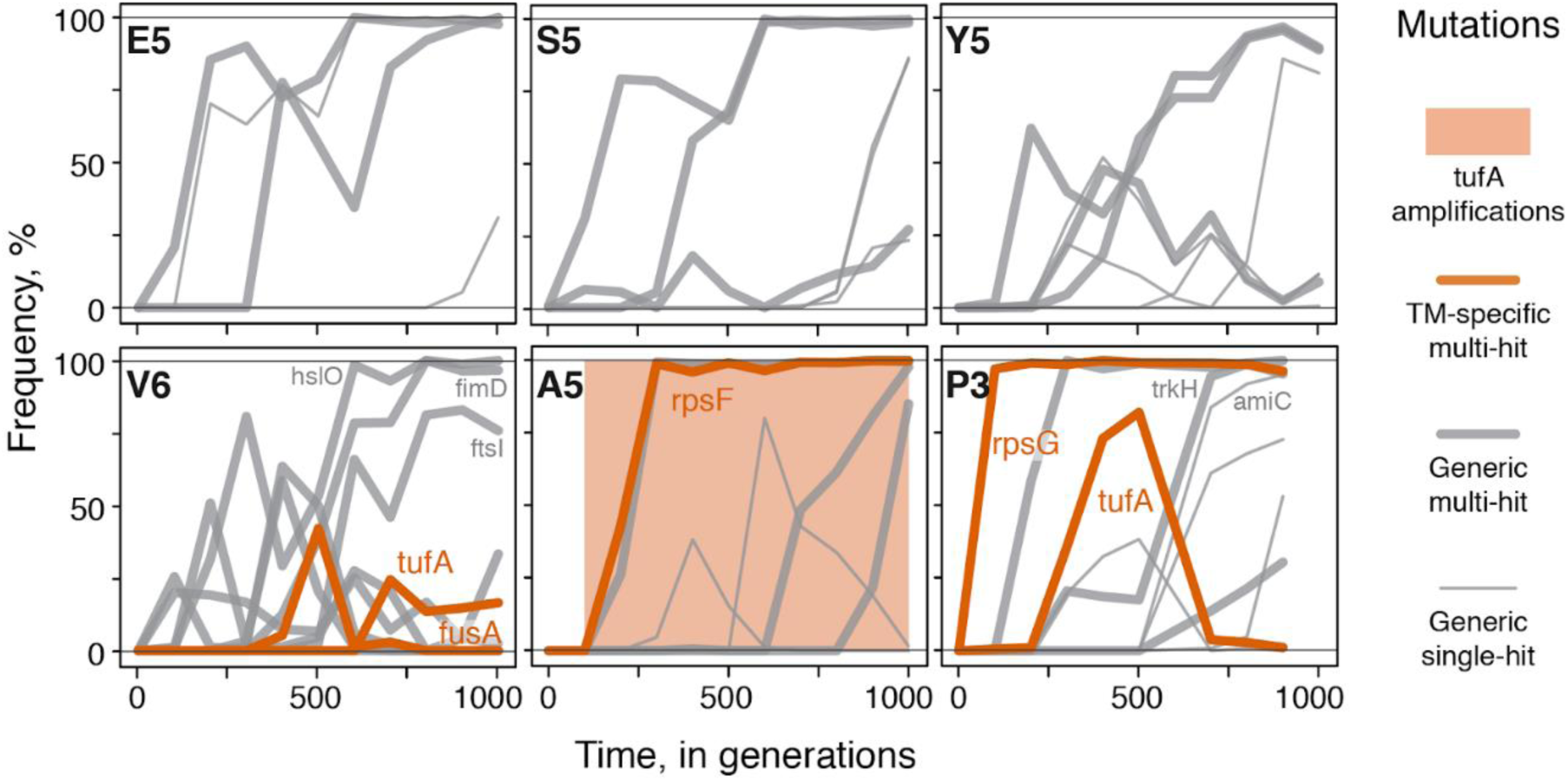
Mutational trajectories in a subset of evolving populations. Mutation frequency trajectories for one representative replicate population per founder is shown (complete data for all sequenced populations can be found in Dataset S6 and SI Appendix, Fig. S3). Each line represents the frequency trajectory of a single mutation. Shading indicates the range of timepoints in which an amplification spanning the tufA locus was detected. Data from generation 1000 in population P3 did not pass our quality control and was removed (Methods). Only mutations mentioned in the text are labelled.

### Evolutionary stalling in the Translational Machinery

Phenotypic evolution of an organism is often interpreted through Fisher’s geometric model (75). In this model, the fitness of a genotype is a decreasing concave function of its distance to a phenotypic optimum. A key property of this model is that genotypes that are farther from the optimum have access to more beneficial mutations with large effects (75). Motivated by Fisher’s model, we hypothesized that TM adaptation is more likely to stall in founders with initially less severe TM-defects because they are presumably closer to their performance optimum. To test this hypothesis, we examined how the TM-specific mutations are distributed among populations derived from different founders.

We found a total of 5, 10, and 12 TM-specific mutations in V, A and P populations, respectively (1.7, 2.0 and 0.8 mutations per population on average, Fig. 3A). We also observed 3, 6 and 2 chromosomal amplifications spanning the *tufA* locus in these populations, respectively. In contrast, we did not observe any TM-specific mutations in the E, S and Y populations (Fig. 3A). The differences in the observed rates of TM-specific mutations between founders are statistically significant (*P* = 4×10^−11^, χ2-test). Consistent with Fisher’s model, the number of TM-specific mutations observed in a founder was negatively correlated with its initial fitness (*r* = –0.993, *P* = 7.8×10^−5^). We also found consistent patterns in the distribution of fixed TM-specific mutations: at least one of them fixed in each of the sequenced A and P populations (Fig. 3A and SI Appendix, Fig. S3), but only 3 out of 6 sequenced V populations fixed one or more TM-specific mutations, assuming that all *tufA* amplifications present in the population at generation 1,000 are fixed (Fig. 3A and SI Appendix, Fig. S3). Thus, in the A and P genetic backgrounds, which have the most severe TM defects, TM-specific beneficial mutations are abundant enough and have large-enough fitness effects that they drive adaptation. In the V founder, which has an intermediate TM defect, TM-specific mutations also contribute to adaptation but to a lesser degree, suggesting that they are less common and/or provide weaker benefits in this background. In the E, S and Y founders whose TMs have only weak defects, TM-specific mutations are either not available or, if they are available, they are so rare and/or so weak that they fail to cross our detection threshold.

**Figure 3.**
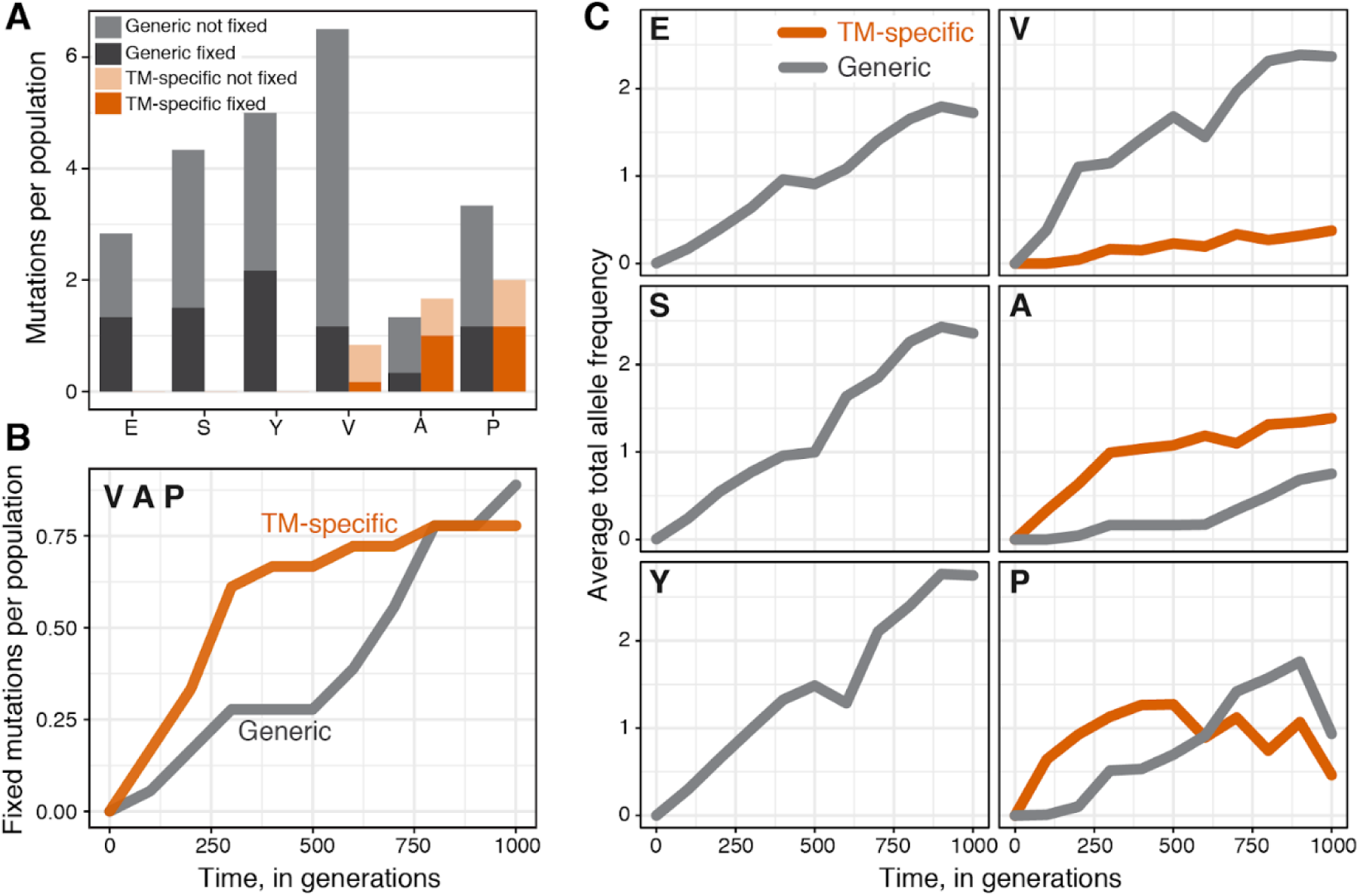
Variation in TM adaptation across founders and time. **A**. Average number of fixed and non-fixed TM-specific and generic mutations per population. **B**. Average number of fixed TM-specific and generic mutations per population over time in the populations derived from V, A and P founders. **C**. Average total allele frequency of TM-specific and generic mutations per population over time. Data in all panels does not include *tufA* amplifications.

If the supply and/or the fitness benefits of TM-specific mutations indeed increase with the magnitude of the TM defect, the contribution of such mutations to adaptation should shrink as populations accumulate TM-specific mutations. To test this hypothesis, we examined the temporal distribution of fixed adaptive mutations in V, A and P populations.

Out of the 14 TM-specific mutations that eventually fixed, 12 (86%) did so in the first selective sweep. As a result, an average TM-specific beneficial mutation reached fixation by generation 300 ± 52, and only one (7%) reached fixation after generation 600 (Fig. 3B and SI Appendix, Fig. S5). In contrast, the average fixation time of generic mutations in the V, A and P populations was 600 ± 72 generations, and 9 of them (56%) fixed after the first selective sweep (Fig. 3B and SI Appendix, Fig. S5). As a consequence, TM-specific mutations were overrepresented among mutations that were fixed within the first 300 generations in the V, A and P populations (*P* = 0.014, Fisher’s exact test). These numbers exclude 11 *tufA* amplifications, for which we do not have temporal resolution. We observe similar trends if we consider the average allele frequencies of mutations (Fig. 3C). We conclude that the accumulation of TM-specific mutations in the A, P and V populations has essentially ceased by the end of the evolution experiment, while the accumulation of adaptive mutations in other modules continued (Figs. 3B, C). In other words, natural selection initially improved both the TM and other cellular modules, but after ∼1 TM-specific mutation was fixed, the focus of selection shifted away from the TM.

Why did TM-specific mutations fix early in evolution? As mentioned above, TM genes occupy only about 4.3% of the *E. coli* genome. It is therefore unlikely that TM-specific mutations arise at higher rates than generic mutations. Instead, their early fixation suggests that their fitness benefits must have been typically much larger than those of generic mutations. To test this hypothesis, we estimated selection coefficients of mutations from the mutation frequency trajectories. We found that mutations in the A and P populations that are detected by generation 300 provide an average selective advantage of 3.7% (*n* = 22), while mutations that are detected after generation 300 provide an average selective advantage of 1.6% (*n* = 28, *P* = 0.0017, t-test). This approach likely under-estimates the effects of the strongest mutations because we cannot accurately resolve selective sweeps that take ≲100 generations. To complement this approach, we genetically reconstructed two TM-specific mutations in their respective founder backgrounds and directly measured their fitness benefits (Methods, Dataset S7 and SI Appendix, Fig. S7). We found that mutation A74G in the *rpsF* gene, which was observed in the population A5, confers a 8.2 ± 0.4% fitness benefit to the A founder. Mutation G331A in *rpsG* gene, which was found in populations P2, P3 and P5, confers a fitness benefit of 6.5 ± 0.5% to the P founder (Dataset S8). Consistent with our hypothesis, these fitness benefits are much larger than the average estimated 1.6% fitness effect of late-arriving mutations (which are mostly generic). Even more strikingly, either of the two reconstructed mutations provides a larger fitness gain to its founder than the total fitness gains achieved by the E, S and Y populations during the entire evolution experiment (Fig. 1A).

Our observations so far can be broadly interpreted in two ways. One explanation is that the TMs in E, S and Y founders are unimprovable by natural selection because they are at their local performance peaks and that TMs in the V, A and P founders become unimprovable after acquiring ∼1 TM-specific mutation. The second interpretation is that natural selection fails to improve the TMs in E, S and Y founders because TM-specific mutations in these backgrounds are too rare and/or too weak to drive adaptation, i.e., TM adaptation is stalled. At the same time, TM adaptation stalls after the fixation of ∼1 TM-specific mutation in the V, A and P populations.

We present three lines of evidence that TM adaptation stalls while the module is still improvable. First, recall that TM-specific mutations fixed in at most three out of six sequenced V populations, which strongly suggests that TMs in at least three remaining V populations (V2, V4, V6) are still improvable by TM-specific mutations (SI Appendix, Fig. S3). Second, we observed that in at least three populations (V5, A6, P3), fixation of one TM-specific mutation was followed by the rise of a second TM-specific mutation, which was eventually outcompeted by a clone carrying generic mutations (see population P3 in Fig. 2 and the other populations in SI Appendix, Fig. S3). This observation suggests that the TMs remained improvable in these populations as well. Third, adaptive mutations have been observed in the translation-related *rpsD* and *rpsE* genes in the long-term evolution experiment in *E. coli* whose founder has a wild-type TM (57). If the wild-type TM is improvable, it is likely that the TM in our E, S, and Y strains—which carry mild defects—are also improvable. It is possible to imagine a structure of epistasis between mutations, such that each of these observations is consistent with the hypothesis that all TMs are at their local performance peaks. However, when considered together, these observations are more parsimoniously explained by the alternative hypothesis: at least some of the TMs in our experiment are not at their local performance peaks; instead, TM-specific mutations fail to keep accumulating in these populations because they are outcompeted by stronger beneficial mutations that improve other cellular modules, i.e., the TM exhibits evolutionary stalling.

Evolutionary stalling prevents a module from reaching its local performance peak and thereby imposes a genetic load, i.e., the organism carrying a stalled module suffers a fitness cost relative to an organism whose module performance is optimal. It is difficult to measure these loads in our populations directly because we do not know how far below their local optima they are. However, we can estimate these loads if we assume that strains with the locally optimal TMs are at least as fit as the E strain. Note that the E strain itself carries a 4% fitness cost relative to the wild-type *E. coli* due to a defect in the TM. 13 out of 20 A and P populations remained significantly less fit than the control E strain, with the average fitness cost of about 0.5% (Fig. 1B). However, TM adaptation in all of these populations stalled before the end of the evolution experiment. Thus, these populations suffer an average genetic load of at least ∼0.5% due to residual TM defects. In fact, the genetic loads could be much higher if the initial TM defects present in the founders have not been alleviated by mutations outside of the canonically annotated TM. Under this assumption, the fact that we did not observe any TM-specific mutations in population V4 would imply that it still carries a ∼19% genetic load imposed by the original TM defect in the V founder. By the same logic, all Y populations still carry a ∼3% genetic load. These estimates suggest that the power of natural selection to improve an individual module can be severely limited by evolutionary stalling.

### Genetic mechanisms underlying evolutionary stalling in the TM

Next, we asked what genetic mechanisms contributed to evolutionary stalling in the TM. We begin with the observation that TM adaptation is stalled or stopped in some founders (E, S, and Y) and initially not stalled in others (V, A and P). This observation indicates that founders have access to different pools of beneficial mutations, i.e., evolutionary stalling is caused by “historical contingency” epistasis (76). To further characterize the extent of historical contingency on evolution in our experiment, we examined the distribution of beneficial TM-specific and generic mutations across populations descended from different founders.

We found that TM-specific mutations were not uniformly distributed among V, A and P populations. Specifically, 4 out of 7 classes of TM-specific mutations arose in a single founder (Fig. 4A). For example, we detected six independent mutations in the *rpsG* gene, which encodes the ribosomal protein S7, and all of these mutations occurred in the P founder (*P* < 10^−4^, randomization test, Methods). Similarly, all four mutations in the *rpsF* gene, which encodes the ribosomal protein S6, occurred in the A founder (*P* < 10^−4^). We have shown above that adaptive alleles at these loci confer strong fitness benefits in their respective founder backgrounds. However, our multiple reconstruction attempts in all other founders were unsuccessful (Methods), which suggests that these mutations are strongly deleterious in other genetic backgrounds, and would explain why we did not observe these mutations in other founders. This result indicates that the severity of the defect in the TM is not the only determinant of the availability of TM-specific beneficial mutations. Instead, defects caused by different foreign *tufA* alleles open up different adaptive pathways within the TM.

**Figure 4.**
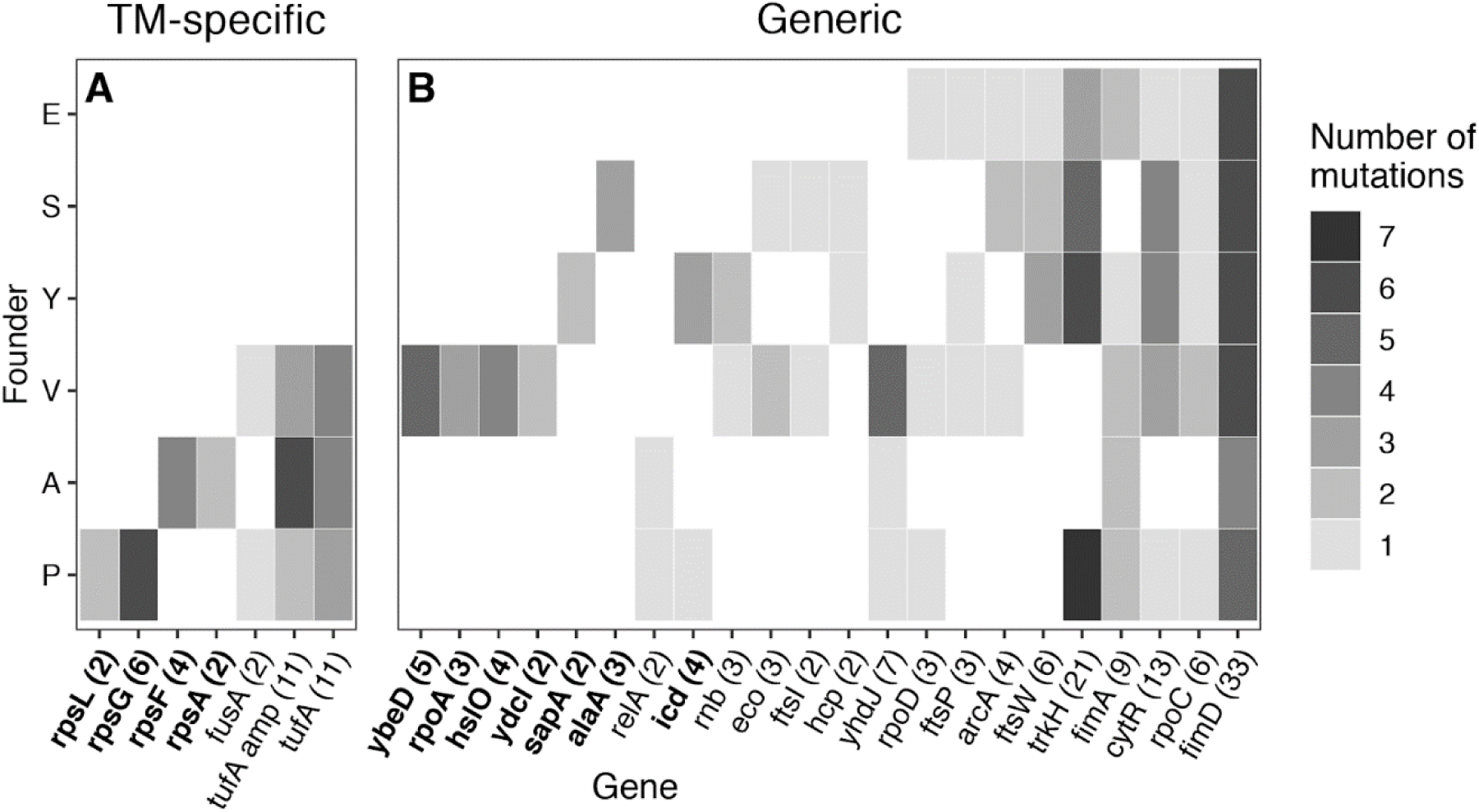
Distribution of putatively adaptive mutations across genetic backgrounds. Heatmap of all putatively adaptive mutations identified via whole-genome sequencing, grouped by founder and by gene. **A**. Translation-associated genes. Amplifications of the *tufA* locus are counted separately from other mutations in *tufA*. **B**. All other genes. Genes in bold are those where mutations were detected in significantly fewer founders than expected by chance (*P* < 0.05, Benjamini-Hochberg correction, see Methods). Numbers in parentheses indicate the total number of mutations in each gene observed across all sequenced populations.

The distribution of some generic mutations among founders was also non-uniform. We found that 7 out of 22 classes of generic mutations occurred in fewer founders than expected by chance (Fig. 4B, Methods). For example, we detected five independent mutations in the *ybeD* gene, which encodes a protein with an unknown function, and all these mutations occurred in the V founder (*P* < 10^−4^). Similarly, all three mutations in the *alaA* gene, which encodes a glutamate-pyruvate aminotransferase, occurred in the A founder (*P* < 10^−4^). To corroborate these statistical observations, we reconstructed the T93G mutation in the *ybeD* gene in all six founder strains and directly measured its fitness effects (Dataset S8). As expected, this mutation confers a 5.9% ± 0.4% fitness benefit in the V founder. In contrast, it is strongly deleterious in the P founder and indistinguishable from neutral in the remaining founders (SI Appendix, Fig. S6). Thus, historical contingency epistasis not only opens up or closes down certain adaptive pathways within the TM but also changes the availability of at least some beneficial mutations outside of it.

We next turn our attention to the genetic mechanisms for the onset of evolutionary stalling in the V, A and P founders. Historical contingency could play a role here too if mutations that accumulate early in evolution alter the identities and fitness effects of subsequent beneficial mutations (40, 57). Additionally, another mechanism, known as “modular epistasis” (40) or “coupon collecting” (57), could also operate if fixation of a mutation depletes the pool of further beneficial mutations in the same module, without changing the effects of mutations in other modules (for example, one beneficial loss-of-function substitution eliminates the benefits of other loss-of-function mutations in the same gene). Modular epistasis could lead to evolutionary stalling in the TM because the most frequent TM-specific mutations with the largest fitness benefits tend to fix earlier thereby enriching the pool of available TM-specific mutations for rare and weak mutations which are more likely to fall below the emergent neutrality threshold (57). If repeated mutations in a given gene occur unusually often in the same population (under-dispersion), this signals historical-contingency epistasis with unknown mutations present in that population (57). On the other hand, unexpectedly even distribution of mutations in a given gene across different populations signals the presence of modular epistasis (57).

To detect historical contingency and modular epistasis, we examined distribution of mutations across populations (*tufA* amplifications were excluded from this analysis; see Methods). We find that TM-specific mutations collectively are overdispersed (*P* = 0.0018). That is, TM-specific mutations at any given locus usually arise only once per population. The distribution of generic mutations collectively is consistent with the random expectation. However, mutations in some individual genes show opposing tendencies. Specifically, mutations in genes *trkH* and *fimD* are overdispersed (corrected *P* < 10^−4^), while mutations in *icd* and *ydcI* are underdispersed (corrected *P* = 0.004 and < 10^−4^). Taken together, these results show that both historical contingency and modular epistasis have both likely contributed to evolutionary stalling in our populations.

## Discussion

The fitness of an organism depends on the performance of many molecular modules inside cells. While natural selection favors genotypes with better-performing modules, it may be difficult for it to improve multiple modules simultaneously, particularly when recombination rates are low and many adaptive mutations in different modules are available. In this regime, natural selection is expected to focus on those modules where large effect mutations occur frequently, while improvements in other modules are expected to stall. Here we have documented such evolutionary stalling in the TM module during laboratory adaptation of *E. coli*.

We found that populations whose TMs were initially mildly perturbed (incurring ≲ 3% fitness cost) adapted by acquiring mutations that did not directly affect the TM. Populations whose TM had a moderately severe defect (incurring ∼19% fitness cost) discovered TM-specific mutations, but clonal interference often prevented their fixation. Populations whose TMs were initially severely perturbed (incurring ∼35% fitness cost) rapidly discovered and fixed TM-specific beneficial mutations. However, after ∼1 TM-specific mutation fixed, further accumulation of mutations in the TM essentially stalled, while mutations in other modules continued to accumulate despite the fact that the TMs remained improvable in at least some of our populations.

We estimated that the genetic load imposed by evolutionary stalling in the TM ranges from 0.5% to 19% in our populations. These estimates are inherently uncertain because we do not know the functional and fitness effects of all fixed mutations. The true genetic loads may be higher than 0.5% because some of the fitness gains in the A and P populations are probably caused by improvements in other cellular modules and because the optimal TM probably provides fitness higher than that of the E strain (since the E strain itself has a TM defect). On the other hand, true genetic loads may be lower than 19% because TM performance may be improved by some mutations in genes not annotated as translation-related.

In our analyses, we defined the TM module operationally as a set of 215 genes in the *E. coli* genome that are annotated as translation-related. However, there may be other genes that contribute to translation that are not annotated as such. A more fundamental problem is that, while the core of the translation machinery is well defined, its periphery may in reality be quite amorphous. In other words, a model of a cell where modules are discrete and clearly delineated is an abstraction. For example, a recent study found that a genetic defect in the DNA replication module can be alleviated by mutations outside of it (54). Similarly, we found that populations with different initial *tufA* defects acquire mutations in statistically distinct sets of genes that are not annotated as translation-related, suggesting that some of these genes may nevertheless be important for translation. Notwithstanding, our results show that the focus of natural selection can transition sharply from one set of genes—those encoding the core of the translation machinery where the initial defect occurred—to functionally more distant parts of the cell such as genes *trkH* and *fimD* that are involved with potassium transport and fimbriae production respectively.

As long as the environment remains constant, the supply of beneficial mutations in an adapting population is gradually being depleted and their fitness effects typically decrease (40, 58–60, 77, 78), thereby lowering the effective neutrality threshold. These changes should in turn allow for less frequent mutations with smaller effects to contribute to adaptation, and adaptation in previously stalled modules may resume. While we did not observe resumption of adaptive evolution in the TM during the duration of this experiment, we find evidence for a transition from stalling to adaptation in *trkH* and *fimD* genes. Mutations in these two genes appear to be beneficial in all our genetic backgrounds (Fig. 4). These mutations are among the earliest to arise and fix in E, S and Y populations where the TM does not adapt (SI Appendix, Figs. S3 and S5). In contrast, mutations in *trkH* and *fimD* arise in A and P populations much later, typically following fixations of TM-specific mutations (SI Appendix, Figs. S3 and S5). In other words, natural selection in these populations is initially largely focused on improving the TM, while adaptation in *trkH* and *fimD* is stalled. After a TM-specific mutation is fixed, the focus of natural selection shifts away from the TM to other modules, including *trkH* and *fimD*.

As the external environment changes, new large-effect adaptive mutations may become available in some modules, which would increase the effective neutrality threshold. This may in turn lead to the onset of evolutionary stalling or prolong the period of stalling in other modules. If environmental fluctuations are sufficiently frequent, e.g., due to seasonality or ecological interactions, and if these fluctuations preferentially open up new adaptive mutations in a subset of cellular modules (e.g., in stress response but not in “housekeeping” modules), some modules may remain stalled for long periods of time despite being improvable, at least in the absence of recombination.

In addition to evolutionary stalling, adaptive evolution in the concurrent mutations regime can have other important consequences for the biology of the organism. For example, Held et al recently showed that fixation of deleterious hitchhiker mutations imposes a limit on the complexity of the organism (37). They also showed that recombination would alleviate this limit. Similarly, recombination is expected to reduce the genetic loads imposed by evolutionary stalling, provided that modules are encoded by tightly linked genes.

Our results give us a glimpse of the fitness landscape of the TM. This landscape appears to be broadly consistent with Fisher’s geometric model (75, 79, 80) in that more defective TMs have access to beneficial mutations with larger fitness benefits than less defective TMs. However, Fisher’s model does not inform us how many distinct genotypes encode highly performing TMs and how they are connected in the genotype space. We observed that the different founders gained distinct TM-specific adaptive mutations. This suggests that the high-performance TMs can be encoded by multiple genotypes that either form a single contiguous neutral network (81) or multiple isolated neutral networks (82). Moreover, we observed that most of our populations with initially severely perturbed TMs were able to discover TM-specific mutations. This suggests that genotypes that encode high-performing TMs may be present in the mutational neighborhoods of many genotypes (81, 83).

In this work, we identified several TM-specific adaptive mutations, but their biochemical and physiological effects are at this point unknown. However, the fact that 11 chromosomal amplifications and 12 noncoding or synonymous events occurred in the *tufA* operon (which consists of *tufA, fusA, rpsG* and *rpsL*) suggests that some of the TM-specific mutations are beneficial because they adjust EF-Tu abundance in the cell. This would be consistent with previous evolution experiments (47, 84, 85). Directly measuring the phenotypic effects of the TM-specific mutations described here is an important avenue for future work.

Overall, our results highlight the fact that it is impossible to fully understand the evolution of a cellular module in isolation from the genome where it is encoded and the population-level processes that govern evolution. The ability of natural selection to improve any one module depends on the population size, the rate of recombination, the supply and the fitness effects of all beneficial mutations in the genome and on how these quantities change as populations adapt. Further theoretical work and empirical measurements integrated across multiple levels of biological organization are required for us to understand adaptive evolution of modular biological systems.

## Materials and Methods

The details of the experimental model, media and growth conditions, fitness and growth measurements, genome sequencing and all statistical analyses are provided in the SI Appendix.

## Data Availability

All strains and plasmids constructed and used in this work are available per request. Raw sequencing data were analyzed with the python-based workflow implemented in Ref. (59) and run on the UCSD TSCC computing cluster via a custom python wrapper script. All analysis and plots reported in this manuscript have been performed using the R computing environment. The script, modified reference genomes and the raw data (except for raw sequencing data) used for analysis can be found at https://github.com/sandeepvenkataram/EvoStalling. Raw sequencing data for this project have been deposited into the NCBI SRA under project PRJNA560969.

## Acknowledgments

We thank members of the Kryazhimskiy and Kaçar groups, Joanna Masel, Ryan Gutenkunst, Suparna Sanyal, Grant Kinsler, Justin Meyer, and Lin Chao for input and feedback. We thank Alex Plesa, Divjot Kaur, Emily Peñaherrera, Kevin Longoria, Lesly Villarejo, Alena Martsul and Sarah Ardell for laboratory assistance. We thank Huanyu Kuo for the analysis of growth-curve data. We thank Eva Garmendia for providing the recombineering plasmids and Georg Rieckh for providing the resistance marker plasmids. We thank Benjamin Good for help with his genome sequencing data analysis pipeline. We thank Kristen Jepsen and the UCSD Institute for Genomic Medicine for sequencing services and the San Diego Supercomputing Center for providing the computational environment. BK acknowledges the support by the John Templeton Foundation (#58562 and #61239); the NASA Exobiology and Evolutionary Biology Program (#H006201406) and the NASA Astrobiology Institute (#NNA17BB05A). SK acknowledges the support by BWF Career Award at the Scientific Interface (#1010719.01), the Alfred P. Sloan Foundation (#FG-2017-9227) and the Hellman Foundation.

## SI Appendix Datasets

**Dataset S1:** Competitive fitness estimates for founder strains relative to the E strain

**Dataset S2:** Competitive fitness estimates for the E founder relative to wild-type *E. coli* MG1655.

**Dataset S3:** Competitive fitness estimates for evolved populations relative to their founder strains.

**Dataset S4:** Competitive fitness estimates for evolved populations relative to the E strain.

**Dataset S5:** Primer names and sequences used for Sanger sequencing validation of 43/45 tested variants.

**Dataset S6:** List of selected (>20% frequency change) variants. Putatively adaptive (locus repeatedly mutated) variants, which were used for all analyses after Figure 2, are annotated in the table.

**Dataset S7:** Primer names and sequences used for strain construction and validation.

**Dataset S8:** Competitive fitness estimates for reconstructed mutants relative to their founder strains.

