## Supplementary material for "Evolutionary Stalling and a Limit on the Power of Natural Selection to Improve a Cellular Module": SI Appendix

### Supplementary Materials and Methods

#### Strains, media and culturing conditions

Liquid medium is the Luria-Bertani medium (LB) (per liter, 10 g NaCl, 5 g yeast extract, and 10 g tryptone) and solid medium is LBA (LB with 1.5% agar), unless noted otherwise. All incubations were done at 37°C, and liquid cultures were shaken at 200 rpm for aeration, unless noted otherwise. All media components and chemicals were purchased from Sigma, unless noted otherwise.

##### Strains and plasmids

All strains in this study were derived from *E. coli* K12 MG1655 as detailed in Ref. (1). Complete methods for the construction of the E, S, Y, V, A and P strains, which harbor a single *tuf* gene variant replacing *tufA* gene, can be found in Ref. (1). Strains with engineered *ybeD*, *rpsF* and *rpsG* mutations were constructed using the same method, except the chromosomal *kanR* marker was not removed (Figure S7). For a full list of primer sequences used for *ybeD*, *rpsF* and *rpsG* engineering, see Dataset S7.

Plasmids pZS1-TnSL and pZS2-TnSL were used in competition assays to provide Ampicillin and Kanamycin resistance, respectively. pZS1-TnSL, derived from pUA66 (2), was kindly provided by Georg Rieckh. pZS2-TnSL was constructed from pZS1-TnSL by replacing the *ampR* cassette with *kanR*.

##### Evolution experiment

Experimental evolution was performed by serial dilution at 37°C in LB broth. To start the evolution experiment, an initial 5 mL overnight culture was inoculated from a single colony from the frozen stock of each founder strains. 10 replicate populations were started from single colonies derived from these overnight cultures. The replicates were serially diluted 1:$10^4$ every 24h (±1h) as follows: 100 µL of saturated culture were transferred into 10 mL saline solution (145 mM NaCl), 50 µL of these dilutions were then transferred to 5 mL fresh LB (tubes were vigorously vortexed prior to pipetting). This resulted in a bottleneck population size of about 5×105 cells. Freezer stocks (200 µL of 20% glycerol + 1 mL saturated culture) were prepared approximately every 100 generations and stored at –80°C.

#### Competitive fitness assays

To carry out pairwise competition assays, an Ampicillin-resistant and a Kanamycin-resistant versions of the query and reference strains/populations were generated by transforming these strains/populations with plasmids pZS1-TnSL and pZS2-TnSL, using standard methods (3). Two replicate competition assays were performed for each query-reference pair with reciprocal markers for our competitions between the evolved populations and their respective founders (four assays total per pair). One replicate for each marker was carried out for competitions between the evolved populations and the E founder. To validate that the resistance-marker plasmids do not differentially impact fitness in any of the six founder genetic backgrounds, we carried out three-way competition assays between the KanR-marked, AmpR-marked and the unmarked versions of the founders (see raw frequency data at https://github.com/sandeepvenkataram/EvoStalling/blob/master/inputData/ThreeWayCompetition_rawFreqData.tab). Since the allele-replacement mutants carry a chromosomal *kanR* marker (see above), they were only competed against AmpR reference strains with 2 replicates.

To start a competition assay, a query and a reference cultures were scraped from frozen stocks and inoculated into 5 mL LB-Amp or LB-Kan media as appropriate. After about 24 hours, the query and the reference cultures were mixed together (in a 1:1 ratio for competitions between the evolved populations and the E founder, and a 1:9 ratio for all other competitions) and diluted 1:10,000 into 5 mL fresh LB media. After that, the mixed culture was propagated as in the evolution experiment. To determine the relative abundances of the query and reference individuals in the mixed culture, 100 µl of appropriately diluted cultures were plated on both LB-Amp and LB-Kan plates after 24, 48 and 74 hours of competition. For some competitions, where fitness differences were particularly large or small, samples from 0 or 96 hours were also obtained. Plates were photographed after a ~24-hour incubation period (when colonies were easily visible) and colonies were automatically counted with the OpenCFU software (4). In each competition, we estimated the fitness of the query strain relative to the reference strain by linear regression of the natural logarithm of the ratio of the query to reference strain dilution-adjusted colony counts against time. Variance was also estimated from these regressions. Replicate measurements were combined into the final estimate using the inverse variance weighting method.

As a concrete example of this analysis, consider the competitions of the founders relative to the E founder. The raw values of ln (query freq / reference freq) at each sampled timepoint for this experiment can be found at https://github.com/sandeepvenkataram/EvoStalling/blob/master/inputData/AncCompetition_lnFreqRatios.tab. For the 4 replicates for the S population, the individual fitness estimates for each replicate are calculated via simple linear regression on this data. The resulting fitness estimates are –4.89E-3, -2.49E-3, -4.81E-3 and -8.88E-3 with variances of 3.79E-4, 7.87E-4, 1.24E-6 and 4.31E-5, respectively. Standard inverse variance weighting allows us to calculate a weighted fitness estimate for this strain (–4.92E-03) and weighted variance (1.20E-06). From here, the standard error can be estimated as well (5.48E-04). The fitness and standard error estimates are converted to percentages (-0.49 ± 0.05%) to generate the final value reported in Table 1.

Competitions between two reciprocally marked versions of the same strain represent a special case. If the two marker-carrying plasmids impose exactly the same fitness cost, our competition assay between two reciprocally marked versions of the same strain is fully symmetric, which implies that in expectation it must yield a fitness value of exactly zero. Any estimate of fitness from a finite number of measurements even in such idealized fully symmetric case will not be zero. However, such deviations from zero would reflect only measurement noise rather than any biologically meaningful fitness difference. In reality, the two marker-carrying plasmids may impose slightly different fitness costs, but because the difference in the cost is undetectable in three-way competitions, we still interpret deviations from zero in our fitness estimates as noise. Therefore, in competitions of reciprocally marked versions of the same strain, we set our fitness estimate to zero and use the four fitness values obtained from the replicate assays to estimate the noise variance as the average of the squared fitness value.

#### Growth rate assays

Strains were inoculated from frozen stock into 5 mL LB media in 15 mL culture tubes and grown overnight. After 24 hours of growth, the cultures were diluted 1:100 into fresh 5 mL of media and grown for 4 hours. They were diluted again 1:100 into 200 µl of LB in flat-bottom Costar 96 well microplates (VWR Catalog #25381-056) and grown in a Molecular Devices SpectraMax i3x Multi-mode microplate reader at 37°C with shaking for 24 hours with absorbance measurements at 600 nm every 15 minutes. Three replicate growth measurements were conducted for each strain. Optical density data were first ln-transformed. A linear regression model was fit to all sets of 5 consecutive data points where OD was below 0.1. Growth rate for the culture was estimated as the maximal slope across all of these 5-point regressions. The mean growth rate and standard error of the mean were calculated from replicate measurements.

#### Genome sequencing

Whole-genome sequencing was conducted for population samples of 6 replicate populations for each of the 6 founders (36 total populations). Each population was sequenced at 11 timepoints, every 100 generations beginning at generation 0. Four lanes of 100 bp paired end sequencing was conducted at the UCSD IGM Genomics center on an Illumina HiSeq 4000 machine. The average per-base-pair coverage across all samples was 131x. Samples E1_t600, E2_t500, Y3_t600, P3_t600, P2_t800, P2_t1000 and A1_t700 yielded data inconsistent with the rest of the allele frequency trajectories from the same population, likely due to mislabelling during sample preparation. These samples were subsequently removed from our analysis.

##### DNA extraction and library preparation

To minimize competitive growth during handling, 100 µl of a 1:10,000 dilution of frozen stock from each sample was plated on LB agar plates and incubated at 37°C overnight. The entire plate of colonies was then scraped and used for genomic DNA extraction. DNA extractions were conducted using the Geneaid Presto mini gDNA Bacteria Kit (#GBB300) following the manufacturer's protocol. Library preparation was conducted using a modified Illumina Nextera protocol as described in Ref. (5).

##### Validation of variants with Sanger sequencing

43/45 variants, particularly those in loci previously annotated to be involved with translation, were validated using Sanger sequencing. Briefly, populations and timepoints containing the variant at substantial frequency were identified, and clones isolated for genomic DNA extraction, PCR and Sanger Sequencing using standard protocols. The primers used for this validation are detailed in Dataset S8. The two mutations that failed to validate were expected to be at relatively low frequency in their populations (17% and 38%), so additional clone sampling may be required to validate these events.

#### Analysis of sequencing data

##### Variant calling

Sequenced samples were mapped to the MG1655 reference genome (NCBI accession U00096.3) and variants were called using a custom breseq-based pipeline described in Supplementary text section 4 of Ref. (6) and kindly provided to us by Dr. Benjamin Good. Briefly, this method leverages the fact that each population was sampled multiple times across the evolution experiment to increase our ability to distinguish real low-frequency variants from sequencing errors and other sources of noise.

The reference genome was modified with the appropriate *tufA* sequence for each genetic background used in the evolution experiment along with the removal of the *tufB* sequence, and annotation coordinates were lifted over to be consistent with the original MG1655 reference sequence using custom scripts. The modified reference genomes and annotation files are included in the github repository. The variants reported in Dataset S5 have been lifted back to be compatible with the original MG1655 reference genome.

##### Annotation

Variant annotation was conducted using the software package ANNOVAR (7). Coding variants were established as normal, while noncoding variants were annotated as being associated with the closest gene (in either strand, in either direction) in the genome, as long as it was less than 1 kb away. As ANNOVAR is not set up to work with *E. coli* by default, the *E. coli* MG1655 nucleotide annotation was downloaded in GFF3 form from NCBI Genbank (U00096.3). Cufflinks (8) gffread tool was used to convert this file to GTF, which was then converted to GenePred by using the UCSC Genome Browser gtfToGenePred tool. The final annotation file was generated using the ANNOVAR retrieve_seq_from_fasta.pl script. The annotation file was lifted over to be compatible with each reference sequence.

Copy number variants were called manually using genome-wide coverage plots generated using samtools (9) "view" command and the R computing environment. As these variants have their frequency confounded with their copy number, only their presence/absence was noted for downstream analysis.

#### Filtering

We considered single nucleotide polymorphisms, short insertion/deletion and the manually identified copy number variants for further analysis. Chromosomal aberrations were ignored because breseq appears to have a high false positive rate (average of 27 "junction" calls per population across all timepoints). Variants were filtered in three successive steps. 1: Variants not identified in multiple consecutive time points were removed. 2: Variants supported by less than 10 reads across all timepoints in a given population were removed. 3: Since we observed fixation events in every population and since there should be no DNA exchange, all truly segregating variants present in a population at generation 100 must either be fixed or lost in generations 900 and 1000. Thus, we removed variants that failed to do so.

Variants that were present at an average frequency ≥ 95% at generation 100 across at least 18 populations were denoted as ancestral mutations that differentiate the founder from the reference genome ($n$ = 10). Variants that were not ancestral but present at ≥ 95% on average across all populations derived from one founder were denoted as founder mutations ($n$ = 11). These mutations were likely introduced as a byproduct of the strain engineering process. Multiallelic variants (two or more derived alleles present in a single population at the same site) were also removed as likely mapping artifacts. Finally, variants that were present at generation 100 in 11+ populations (of 36 total sequenced populations) are either mapping artifacts or pre-existing variants and were not considered further ($n$ = 169, including the 10 ancestral mutations identified earlier).

#### Identification of adaptive mutations

The putatively adaptive mutations were identified as follows. We first identified mutations that reached at least 10% frequency, were present in at least two consecutive time points and whose frequency changed by at least 20% throughout the evolution. We then merged together such mutations within 10 bp of each other as likely being derived from a single event. This resulted in a set of candidate adaptive mutations. To identify likely adaptive mutations in this candidate set, we considered only mutations in "multihit" genes, i.e., genes with 2 or more candidate adaptive mutations.

#### Identification of modules in the genome

The 215 genes annotated as being associated with translation were identified using the Gene Ontology database at http://geneontology.org/ by searching for all *E. coli* K12 genes that were identified in a search for "translation OR ribosome".

#### Estimation of selection coefficients from frequency trajectory data

Selection coefficients were estimated independently for every putatively adaptive mutation. The allele frequency trajectory of each such mutation was cleaned to remove all timepoints where the frequency was either 0, 1 or not determined due to a missing datapoint. We then consider the subset of these points where the frequency of the mutation consistently increased by at least 5% between consecutive values, regardless of if those consecutive values represent consecutive generations or not. We then perform essentially the same calculation as in our fitness assays above. Assuming that all other individuals in the population that are not carrying this mutation constitute a monomorphic "reference strain", we calculate log(mutation frequency / (1 - mutation frequency)) for our log frequency ratio. We then compute the slope of these estimates vs the generations in which they were sampled to estimate the fitness benefit of each mutation. Note that if multiple adaptive mutations increase in frequency simultaneously as part of a single cohort (i.e. they are genetically linked), each such mutation will still have their fitness benefit calculated independently.

#### Statistical analyses

The expected number of mutations in multi-hit genes was calculated via multinomial sampling. Mutations were randomly redistributed across all genes in the *E. coli* genome controlling for variation in gene length. The average of 10,000 such randomizations was used to calculate an empirical FDR. A similar procedure was used to estimate the probability of observed as many or more TM-specific mutations by chance as we actually observed in this study.

To test whether mutations in the 7 TM-specific multi-hit loci were distributed uniformly across the six founders we first estimated the entropy of the distribution of mutations across founders for each gene. Mutations in that gene were then randomly redistributed across six founders 10,000 times, weighted by the total number of TM-specific mutations observed in each founder. An empirical *P*-value was calculated as the fraction trials with smaller than observed entropy value. These *P*-values were then corrected for multiple testing across the 7 TM-specific loci using the Benjamini-Hochberg procedure. We used the same procedure to test for significant deviations in the distributions of generic mutations across founders.

We then tested whether mutations at each multi-hit locus were over- or under-dispersed across populations via permutation analysis. First, we identified all founder backgrounds in which mutations at a given locus were observed. We then calculated the entropy of the distribution of these mutations across these populations as our test statistic. We then resampled the same number of mutations 10,000 times uniformly across these populations and obtained the null distribution of the entropy statistic. We calculated an empirical P-value for the observed entropy, testing separately for overdispersion (observed entropy is significantly higher than expected) and underdispersion (observed entropy is significantly lower than expected). We also calculated whether TM-specific mutations were collectively over- or under-dispersed by summing the entropies of individual TM-genes. We conducted a similar analysis for generic loci.

#### Supplemental Figures

**Figure S1.** Growth rate of founders vs fitness relative to the E strain.

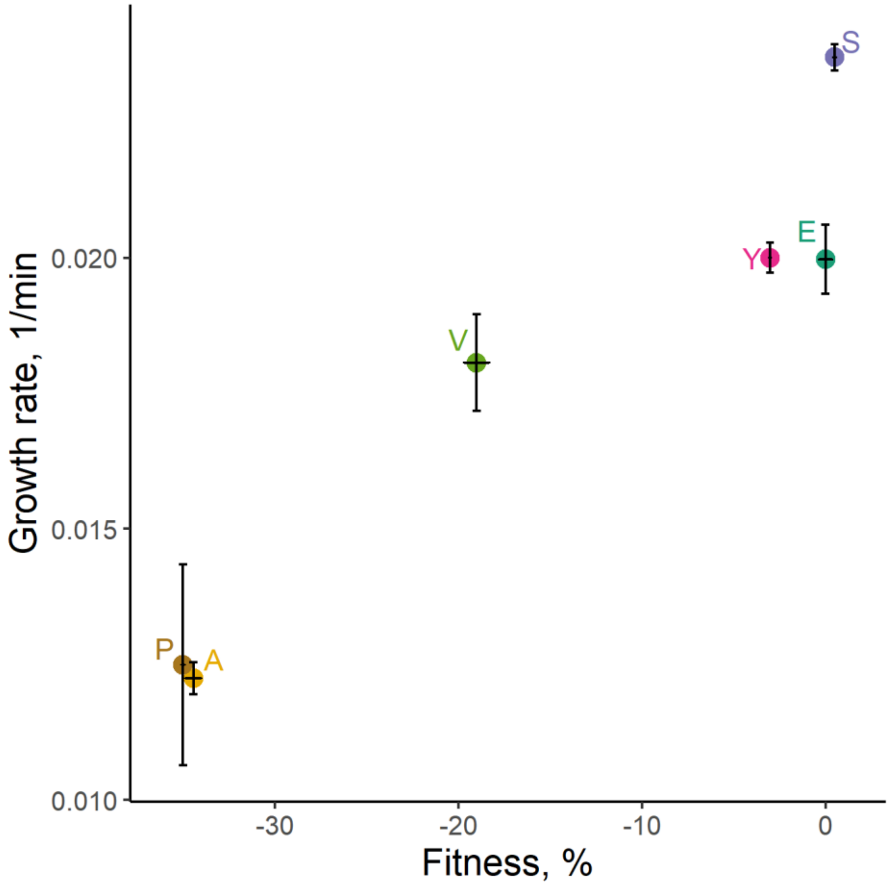

**Figure S2.** Genome-wide coverage of 11 populations with evidence of *tufA* amplifications shown at one timepoint where the amplification is detected. The red vertical dashed lines denote the genomic coordinates ~50kb upstream and downstream of the *tufA* locus.

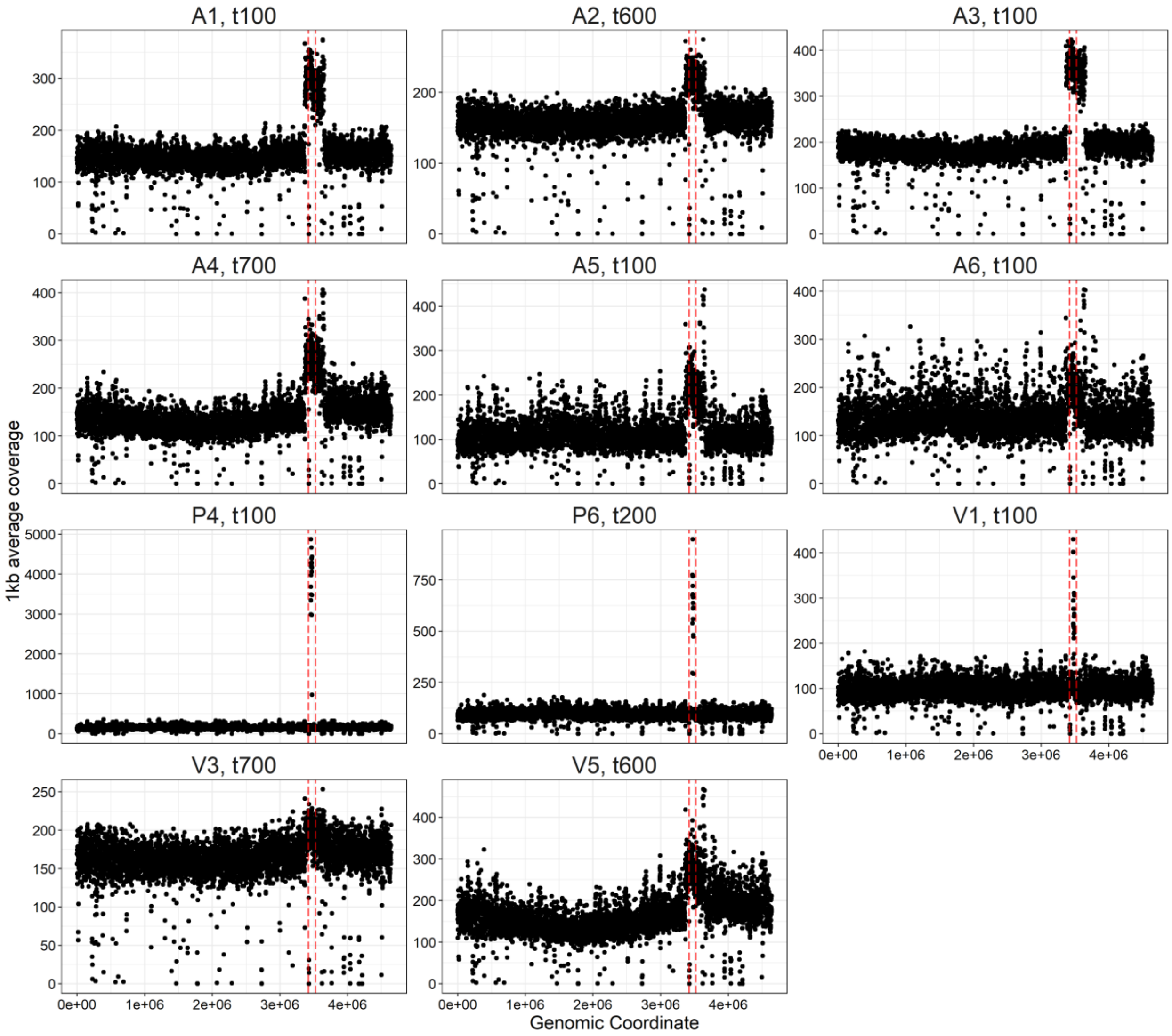

**Figure S3.** Mutational trajectories for all 36 sequenced populations. Each line represents a single mutation that changed in frequency by > 20%. Thin lines represent mutations in loci that were only mutated once across the entire dataset, while thick lines are loci that were multiply mutated. Orange shading indicates timepoints in which an amplification spanning the *tufA* locus was detected in that population as in Figure 2.

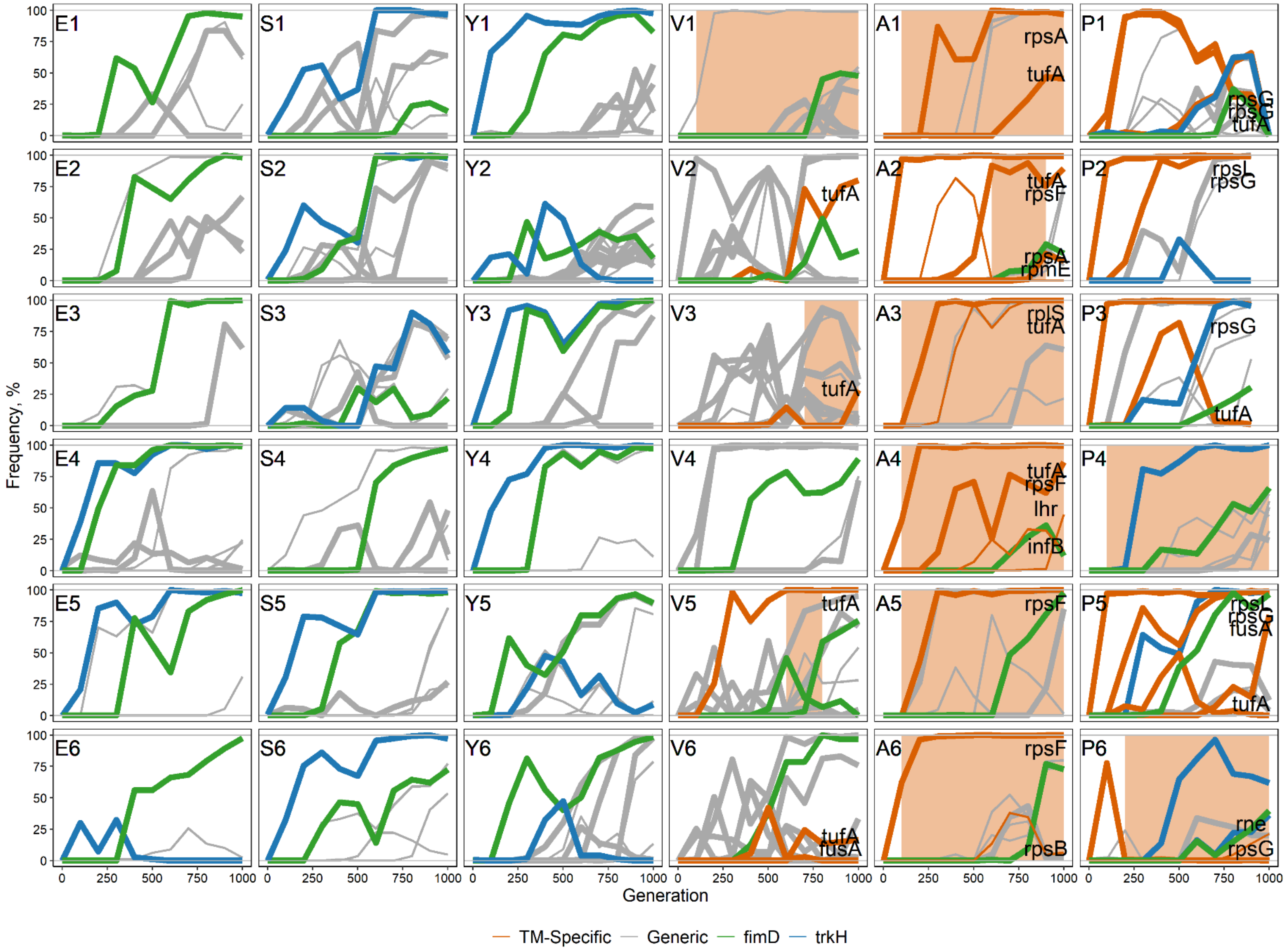

**Figure S4.** Similar to SI Appendix Figure S3 but for mutations with a relaxed criterion for frequency changes (> 10% instead of > 20%). This relaxation of our filtering criteria increases the total number of detected selected mutations from 288 to 423 and the number of putatively adaptive mutations in multi-hit genes from 178 to 263. The FDR for putatively adaptive mutations in multi-hit genes is 21.9%. 6 new TM-specific mutations appear in this figure compared to SI Appendix Figure S3, but the number of multi-hit TM genes remains unchanged.

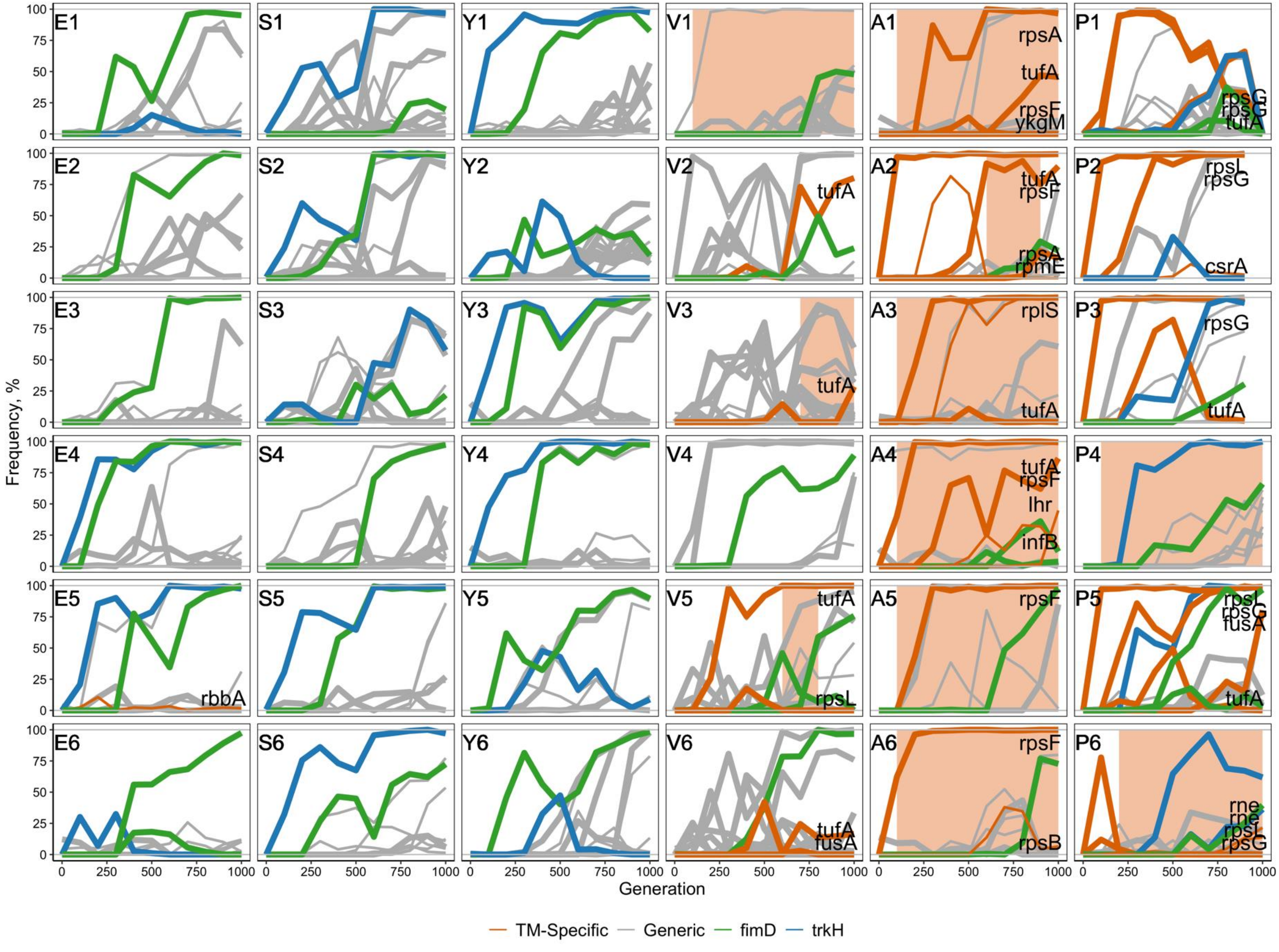

**Figure S5.** Similar to Figure 3B but shows the average number of fixed mutations in each module for each founder separately rather than in bulk. *trkH* and *fimD* mutations are shown as sub-groups of the Generic module.

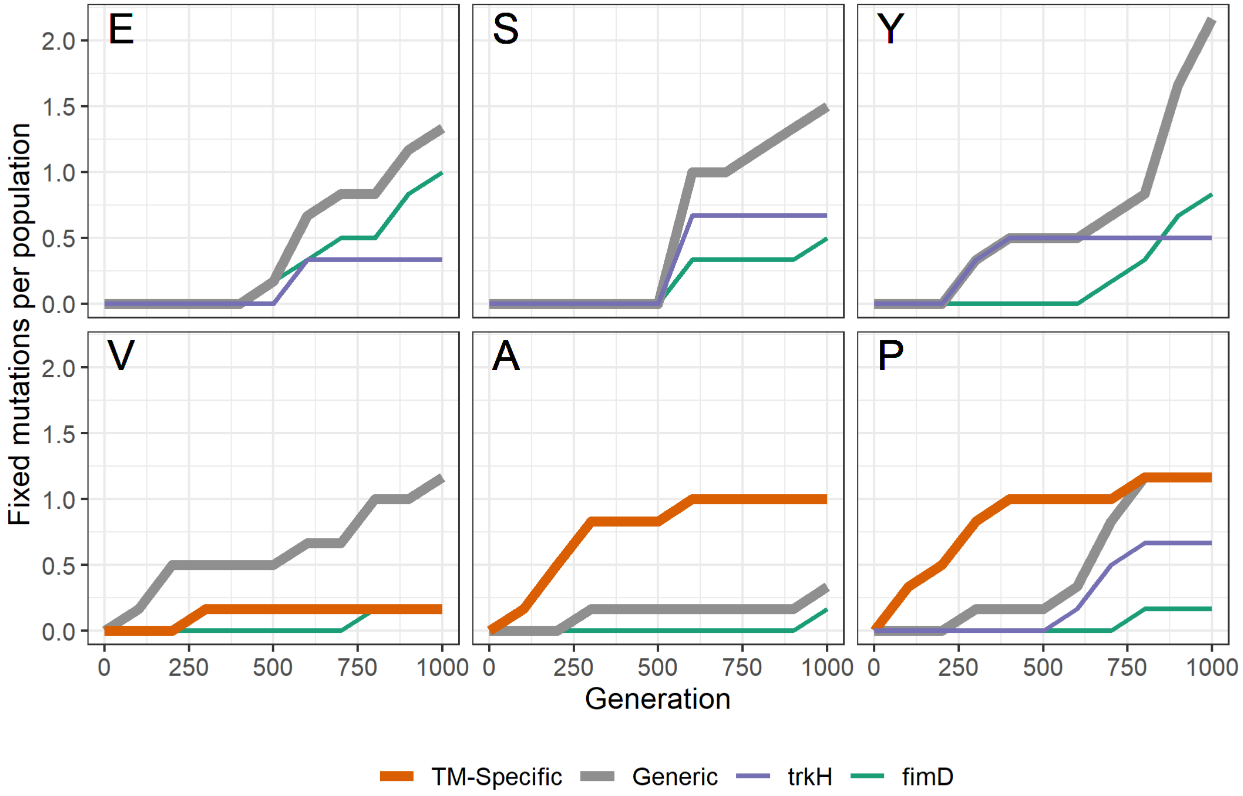

**Figure S6.** Fitness effect of the *ybeD* T193D mutation in different founders. Fitness effect of the mutation is measured in a direct competition of each founder with the mutation against the founder without the mutation. Error bars show ±1 SEM.

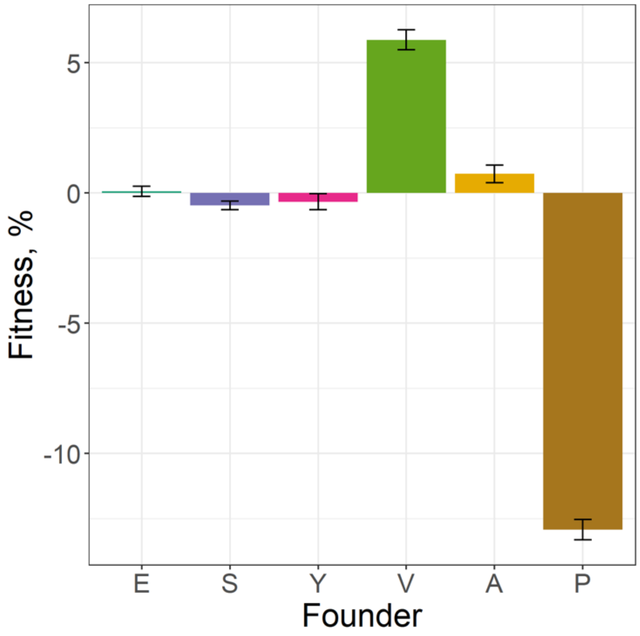

**Figure S7.** Methods for integrating mutations into the genome.

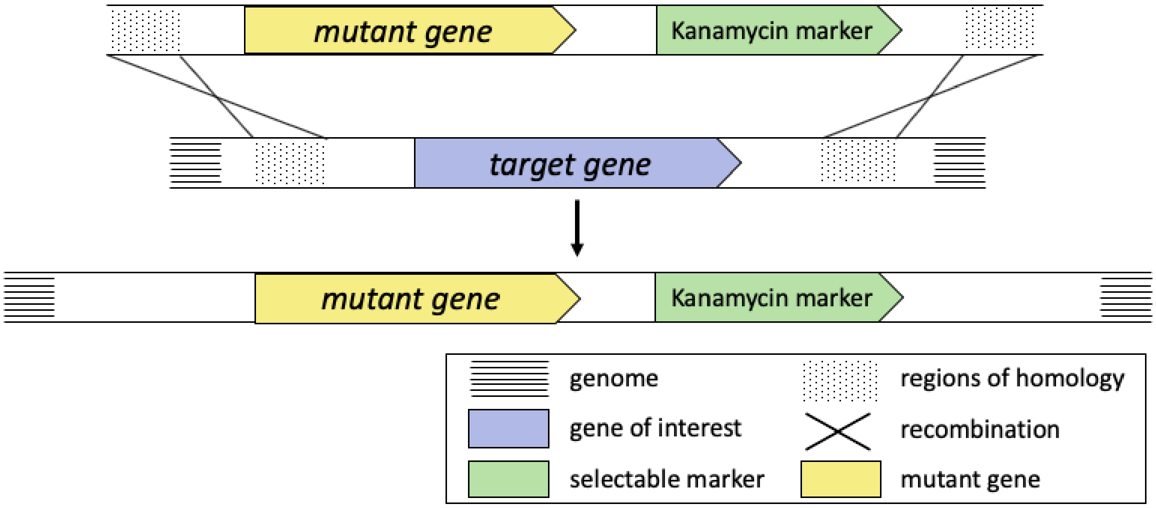
